# Visual Long-term Memory Can Replace Active Maintenance in Visual Working Memory

**DOI:** 10.1101/381848

**Authors:** Mark W. Schurgin, Corbin A. Cunningham, Howard E. Egeth, Timothy F. Brady

## Abstract

Humans have remarkable visual long-term memory abilities, capable of storing thousands of objects with significant detail. However, it remains unknown how such memory is utilized during the short-term maintenance of information. Specifically, if people have a previously encoded memory for an item, how does this affect subsequent working memory for that same item? Here, we demonstrate people can quickly and accurately make use of visual long-term memories and therefore maintain less perceptual information actively in working memory. We assessed how much perceptual information is actively maintained in working memory by measuring neural activity during the delay period of a working memory task using electroencephalography. We find that despite maintaining less perceptual information in working memory when long-term memory representations are available, there is no decrement in memory performance. This suggests under certain circumstances people can dynamically disengage working memory maintenance and instead use long-term memories when available. However, this does not mean participants always utilize long-term memory. In a follow-up experiment, we introduced additional perceptual interference into working memory and found participants actively maintained items in working memory even when they had existing long-term memories available. These results clarify the kinds of conditions under which long-term and working memory operate. Specifically, working memory is engaged when new information is encountered or perceptual interference is high. Visual long-term memory may otherwise be rapidly accessed and utilized in lieu of active perceptual maintenance. These data demonstrate the interactions between working memory and long-term memory are more dynamic and fluid than previously thought.

## Introduction

Visual working memory is an online system used to actively retain and manipulate information over brief periods (Baddeley & Hitch, 1974; Vogel, Woodman & Luck, 2006; Cowan, 2008; Ma, Husain & Bays, 2014), acting as bridge across spatial and temporal gaps in our perception (Cowan, 2008; Hollingworth, Richard & Luck, 2008; Hollingworth, Matsukura & Luck, 2013). The most notable characteristic of visual working memory is that it is capacity limited (Vogel, Woodman & Luck, 2001; Awh, Barton & Vogel, 2007), with individual capacity differences strongly correlating with measures of broad cognitive function, including fluid intelligence and academic performance (Alloway & Alloway, 2010; Fukuda et al., 2013). By contrast, visual long-term memory is the storage of visual information operating through the retrieval of memory traces over any time scale without continued active maintenance (Squire, 2004; Cowan, 2008; Brady, Konkle & Alvarez, 2011). A large body of work has demonstrated humans have remarkable visual long-term memory abilities (Shepard, 1967; Standing, 1973; Biederman, 1987), capable of storing thousands of objects with significant detail (Brady, Konkle, Alvarez, Oliva, 2008).

In order to isolate these two systems and study visual working memory without contributions from long-term memory, visual working memory is often studied using simple displays consisting of stimuli such as meaningless colored squares or line orientations. However, in the real-world visual working memory does not operate in isolation, and in our everyday life we rarely need to hold in mind semantically meaningless stimuli or stimuli that we have never previously encountered. Existing work has shown the important role of semantic meaning and expertise for visual working memory. For example, individuals have greater working memory capacity for kinds of objects for which they have more expertise (e.g., faces; Curby, Glazek & Gauthier, 2009) and having relevant semantic knowledge – e.g., remembering a mug rather than a colored circle – also results in improvements in the capacity of visual working memory (Brady, Stormer & Alvarez, 2016; O’Donnell, Clement & Brockmole, in press).

However, the role of previous encounters that result in visual long-term memory traces for a *specific item* have rarely been investigated, and those investigations have focused on perceptually simple, meaningless stimuli (e.g., Carlisle et al. 2011). By long-term memory traces, we are not referring to semantic or categorical knowledge about previously seen items. Rather, we are referring to specific visual long-term memory representations – memories of having seen a particular object – that may be utilized to inform performance. Imagine a friend asks you to pour them some water and points to a unique red cup with a specific design to indicate it is theirs. To complete this task, you need to hold in mind which cup is theirs while you fetch the water. But what if an hour earlier you had encoded a memory that your friend’s cup was this particular design of red cup? Does this existing long-term memory reduce or eliminate the need to hold this perceptual information about your friend’s cup actively in mind, thus alleviating demands on the capacity-limited visual working memory system? Or when needing to act on information in the immediate future, do people always use both systems, resulting in a reduced chance of making an error but coming at a cognitive cost of having to engage active maintenance of perceptual information in working memory even when an existing memory could serve to guide their performance?

Surprisingly, this question remains largely unexplored. The majority of research investigating the connection between working memory and long-term memory focuses on the ways in which working memory acts as a passageway through which information must pass at it progresses onward to long-term memory (Cowan, 2008; Schurgin, 2018), or the extent to which representations that are held active in working memory may be just activated versions of underlying long-term memories with similar representational content (e.g., Brady et al. 2013) with working memory capacity constraints arising from keeping these representations activated (Cowan, 1999; D’Esposito & Postle, 2015). One of the main constraints on participants utilizing existing long-term memory representations in lieu of engaging working memory is that participants would have to almost immediately encode an item differentially depending upon whether an item was previously encoded; any delay would significantly slow down task performance, or, in an experimental situation, cause them to miss the object entirely. There is some existing evidence that participants can at least under some conditions quickly and accurately access existing long-term memories to perform tasks, such as utilizing long-term memory systems to efficiently search for targets in a display or to answer questions related to their own expertise (e.g., Ericsson & Kintsch, 1995; Wolfe, 2012; Cunningham & Wolfe, 2014). In addition, some evidence has shown that when a small set of simple stimuli are repeatedly required for an attentional task, they may no longer be held actively in working memory during a delay, but may instead be offloaded to long-term memory (Carlisle et al. 2011). Thus, it is possible that under some conditions participants would be able to access passively stored visual long-term memories efficiently enough to avoid having to engage working memory when they see the items and form a new, activated representation of those items.

In the present study, we used electrophysiological recordings to directly test whether or not having a previous long-term memory of an item alters or eliminates the need for active maintenance for that same item in working memory. One particularly strong marker of active maintenance in working memory is the contralateral delay activity (CDA), a sustained negative activity in the posterior region of the brain that occurs contralateral to a set of lateralized objects that are being remembered over a delay period (Klaver, Talsma, Wijers, Heinze, & Mulder, 1999; Vogel and Machizawa, 2004). CDA amplitude increases when more items are stored in working memory, and decreases immediately when items are dropped from working memory (Vogel et al., 2005). It increases when more items are remembered in a way that correlates with individual working memory capacity (Vogel and Machizawa, 2004). And it’s correlated with behavior, such that it’s larger on correct than incorrect trials (McCollough et al., 2007). Thus, the CDA provides a useful measure to assess whether and how long-term memory for a specific item may affect the active maintenance of perceptual information in visual working memory, since it can be utilized to assess how much information is being held actively in mind.

In Experiment 1, participants completed a working memory task, holding two images of real-world objects in mind, followed by a perceptual discrimination test focused on one of those objects. For half the trials, both of the objects that needed to be remembered were completely new, and for the other half, one of the two objects had been previously encountered during a long-term memory encoding session. Using the CDA as an index of visual working memory maintenance activity, we tested whether participants could quickly access their existing long-term memory representations in order to avoid having to hold previously encountered items in visual working memory. We found that, indeed, participants could use long-term memory to reduce or replace the need to hold items actively in mind with no cost to behavioral performance, suggesting extremely quick access to visual long-term memories and a fluid relationship between visual long-term memory and active maintenance in visual working memory. In Experiment 2, we explored what conditions would make active maintenance necessary even when an existing long-term memory was available. We hypothesized that active maintenance may be used when long-term memories are unreliable. Interference is a well-known issue for long-term memory, and is theorized to be one of the primary reasons for having a separate working memory system (e.g., Engle, 2002; visual working memory is robustly tolerant to interference, see Schurgin & Flombaum, 2018). Accordingly, we significantly increased perceptual interference in the working memory task to make the usage of long-term memory less reliable. Under this circumstance, we did not find any differences across conditions in CDA amplitude. Together, our data suggest that when perceptual interference is low, participants are able to quickly and accurately use long-term memory in lieu of working memory, reducing the cognitive demands on working memory by requiring less perceptual information to be held in mind.

### Experiment 1: Visual Long-Term Memory Replaces Active Maintenance in Working Memory

To assess the potential interaction of long-term memory and active maintenance in visual working memory, in Experiment 1 participants completed a working memory task, holding two images of real-world objects presented sequentially in mind, followed by a perceptual discrimination test focused on one of those objects. For half the trials, all the objects encountered were completely new, and for the other half both of, one of the two objects had been previously encountered during a previous episodic memory encoding session. Critically, the experiment was designed with two key components: (1) during the working memory task participants had to make a perceptual discrimination at test, which means semantic, categorical or gist information was insufficient to give the correct answer. Thus, if observers utilized information from long-term memory at test, visual details in these representations would be necessary to inform performance. (2) Only half the trials contained an item that was previously encoded in long-term memory, during these trials the long-term item could appear either as the first or second item in the sequential presentation, and at subsequent test there was only a 50% chance the previously encoded long-term memory item would be tested. As a result, participants could not expect whether or not a previously encoded long-term memory item would be present on a trial, and if it did appear whether or not it would be subsequently tested, in order to preemptively disengage working memory resources.

## Methods

### Participants

23 Johns Hopkins University undergraduates participated in Experiment 1. The results from three participants were excluded due to technical errors and/or excessive artifacts in the EEG signal, leading to a final sample of 20 participants. All participants reported normal or corrected-to-normal visual acuity. Participation was voluntary, and in exchange for extra credit in related courses. The experimental protocol was approved by the Johns Hopkins University IRB.

### Procedure

Experiment 1 began with a long-term memory encoding task (Figure 1A). Participants viewed 120 images of real-world objects (stimuli taken from Schurgin & Flombaum, 2017), one after another for 2 seconds each (500 ms ISI), and were explicitly told to remember each image as best they could. After completing the task, approximately 30 minutes later participant then completed a lateralized visual working memory task.

**Figure 1.**
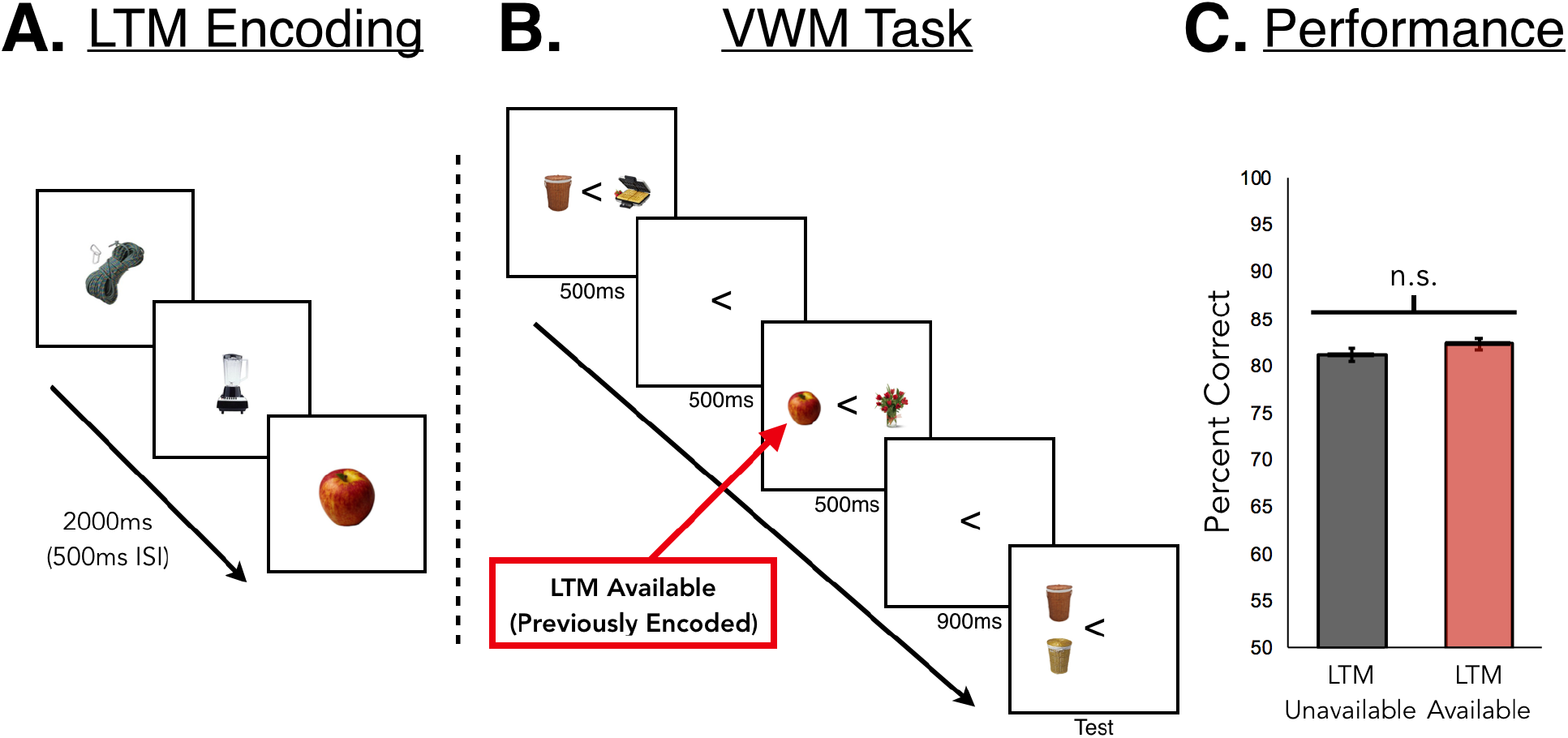
(**A**) Participants first completed a long-term memory encoding session. (**B**) After the long-term memory encoding session and a delay, participants completed a sequential working memory task. To allow measurement of the contralateral delay activity, a neural marker of working memory, participants were cued to remember only the objects on either the left or right side of fixation. For half the trials, both studied images had never been previously encountered (LTM-unavailable condition). For the other half of trials, one of the images had been seen previously in during an episodic long-term memory encoding session (LTM-available condition, shown above). Objects were presented sequentially for 500ms each, with a 500ms ISI. After a 900ms delay, a perceptual discrimination 2-AFC test assessed detailed object memory. (**C**) Behavioral results for Experiment 1. We found no difference in performance based on whether observers had a previous long-term memory representation available or not. Error bars represent within-subject standard error (Cousineau, 2005).

Participants completed 240 total trials of the working memory task. On each trial, four categorically distinct objects were presented. Objects were presented in pairs sequentially (500 ms each, 500 ms ISI) in each visual hemifield. Participants were cued at the beginning of the trial to remember the objects only on either the left or right of fixation and to ignore the other objects. Thus, during a given trial, participants needed to only remember two categorically distinct objects (presented on either the left or right side of the screen). After both pairs of objects were presented, this was followed by a delay interval (900 ms) and then a memory test (Figure 1B). As a result of presenting pairs of objects sequentially during encoding, the visual display on both hemifields were equated for perceptual information so that any brain differences between the cued and uncued side during the delay interval were due to differences in holding the items in working memory. Objects on the to-be-ignored side of the display were drawn from a separate set of categorically distinct real-world objects from the same stimulus set, which were never shown during long-term memory encoding or on the to-be-remembered side of the working memory display.

The cue indicating which side of the screen to remember was an arrow pointing either left or right that appeared at the center of the screen 1,000 ms before the presentation of the objects. Participants were instructed to keep their eyes in the center of the screen throughout each trial until the test display appeared. Trials with horizontal eye movements were excluded from the analysis. On half of the trials both of the to-be-remembered objects presented were new (i.e. never encountered during the previous long-term memory encoding session), which we term “LTM-unavailable” trials. These trials provided a baseline assessment of working memory performance in the task. For the other half of the trials, one of the objects encountered had been previously encoded during the long-term memory encoding session, which we term “LTM-available” trials. Of these trials, half of the time the long-term memory item was encountered as the first item in the trial sequence (“LTM-first”), whereas for the other half the long-term memory item was encountered as the second item in the sequence (“LTM-second”).

After the 900-ms delay interval, participants were presented a perceptual discrimination forced-choice test that remained visible until participants made a response. This test consisted of one of the two previously studied items from the working memory display (counterbalanced equally by presentation order across all conditions) and a second, similar-looking lure, with both items appearing above or below fixation on the to-be-remembered side. Participants were instructed to indicate which of the two objects had appeared in the previous display. Since the test included a previous item from the working memory display and a similar-looking lure, participants could not rely on gist, categorical, or semantic information in order to identify the correct item. Rather, visual details in memory were necessary at test.

Importantly, in this design, even if a participant recognized a previously encoded long-term memory object during the working memory task, this was not indicative of which item they would be tested on. In addition, the sequential presentation design ensured that participants could not direct attention solely to the new item, as they would be able to do if the items were presented simultaneously. The sequential nature of the task also allowed us to examine potential distinction between gating and maintenance in working memory (Badre, 2012); in particular we could examine whether once working memory was engaged (i.e., when encountering a new item first in a sequence), people tended to actively maintain the second item as well (even when this item had an existing long-term memory available), or whether each item was treated independently.

### Electrophysiological Recordings and Analysis

EEG was recorded continuously from 32 Ag/AgCL electrodes mounted in an elastic cap and amplified by an ActiCHamp amplifier (BrainVision). EEG data were sampled at 500 Hz. Data were online referenced to Cz and later were offline referenced to the average of the right and left mastoid. Eye movements were measured through the two frontal eye channels (FP1 and FP2).

Continuous EEG data were filtered offline with a band pass of 0.01–112 Hz. Trials with horizontal eye movements, blinks, or excessive muscle movements were excluded from the analysis. All EEG and ERP data analyses were conducted through EEGLAB (Delorme & Makeig, 2004) and ERPLAB (Lopez-Calderon & Luck, 2014) toolboxes for Matlab. Timing of the stimulus presentation and event codes were monitored and corrected using a photodiode. ERPs were time-locked to the onset of the memory display in all experiments, and ERPs from artifact-free epochs were averaged and digitally low- pass-filtered (−3-dB cutoff at 25 Hz) separately for each subject.

ERPs elicited by the memory display were averaged separately for each condition and were then collapsed across tø-be-remembered hemifield (left or right) and lateral position of the electrodes (left or right) to obtain waveforms recorded contralaterally and ipsilaterally to the to-be-remembered side. For each participant, mean CDA amplitudes were measured with respect to a 200-ms prestimulus period at four lateralized posterior electrodes (PO3/PO4/PO7/PO8), consistent with existing data on the location of the CDA (McCollough, Machizawa & Vogel, 2007).

The measurement window for the CDA started 300 ms after the offset of the memory display and lasted for 400 ms, until the cue indicating the memory test item. Thus, the CDA amplitude was measured between 1800-2200 ms with respect to the onset of the memory display. The resulting mean amplitudes were statistically compared using paired t-tests and ANOVAs.

## Results

We first examined participants’ accuracy separately in the LTM-available and LTM-unavailable conditions. We found that across both conditions, accuracy was similar. In particular, participants averaged 81.2% correct (SEM = 1.6%) in the LTM-unavailable condition and 82.4% correct (SEM = 1.4%) in the LTM-available condition, which were not significantly different from one another, t(19) = 0.92, p = 0.37, Cohen’s dz = 0.20 (Figure 1C). Furthermore, we found no effect of order in the LTM-available condition. Participants averaged 81.2% correct (SEM = 1.7%) in the LTM-first condition and 83.6% correct (SEM = 1.5%) in the LTM-second condition, t(19) = 1.46, p = 0.16, Cohen’s dz = 0.32. Additionally, in all the LTM-available conditions we found no performance difference for whether the old or new item was tested (all p’s > 0.2). Thus, having previously encoded objects in long-term memory didn’t offer any advantage or disadvantage in performance over maintaining new objects in working memory. This is consistent with either participants not utilizing long-term memory at all in the task, or with the idea that participants are strategically making use of active maintenance in working memory only when they do not have a strong long-term memory representation available. However, this does provide evidence that participants do not use the two kinds of representations additively.

We next assessed CDA activity, our neural marker of how much information was being actively maintained in working memory. Due to the short time period between the two sequentially presented stimuli and the predictable timing (and thus possibility of preparatory activity), we focused only on the longer 900ms delay interval after the second object was presented and before the test (We used sequential presentations largely to prevent people from distributing attention to only the new items). First, we compared trials containing only new images, our working memory baseline (LTM-unavailable), to trials containing a previously seen image (LTM-available). Consistent with the hypothesis that long-term memory was being used to substitute for active maintenance in working memory, we found that the CDA amplitude was lower when participants encountered a previously seen object (M = 1.20, SEM = 0.21) compared to when both images were new (M = 1.63, SEM = 0.19), t(19) = 2.17, p = 0.04. Combined with the lack of behavioral advantage for the LTM-available condition, this provides evidence that participants used visual long-term memories in lieu of holding items active in working memory. In particular, participants seem to be using visual long-term memory to substitute for or replace active maintenance in working memory when they encounter a familiar item.

**Figure 2.**
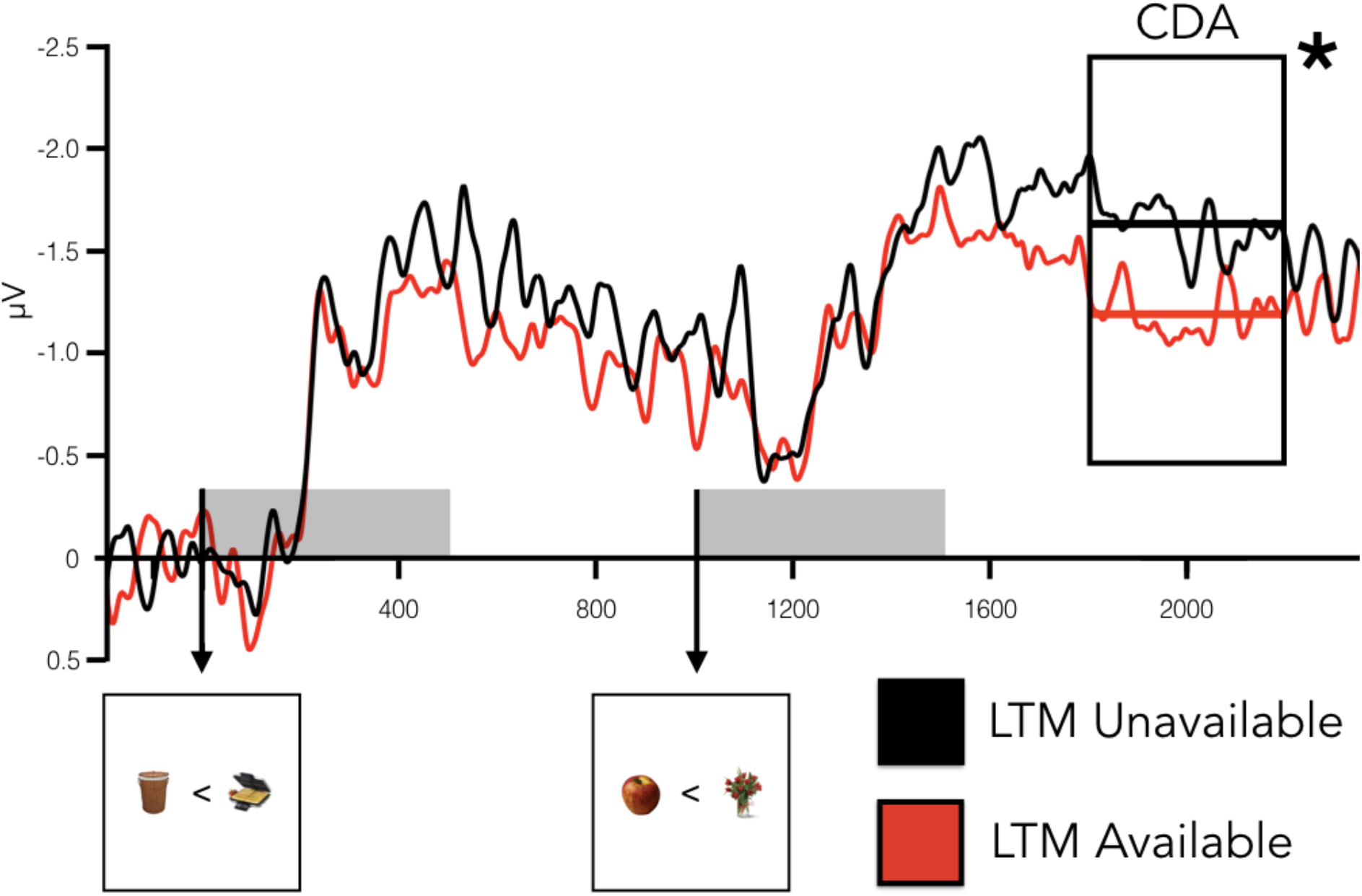
Results of Experiment 1. Contralateral-minus-ipsilateral waveforms for the LTM Unavailable (black) and LTM Available (red) conditions. The CDA is measured from 300 ms after offset during the delay period (black rectangle, labeled CDA). We observed significantly reduced CDA amplitudes for when participants had a LTM representation available, compared to when they did not.

The design of our experiment, with sequential presentations and thus LTM-first and LTM-second trials, also allows us to examine the role of "gating" in the engagement or lack of engagement of visual working memory. A significant literature has suggested that working memory operates via separate gate and maintenance mechanisms: when the gate is open available information can enter working memory, and when the gate is closed the contents of working memory are sustained while keeping irrelevant information out (Miller & Cohen, 2001; Raghavachari et al., 2001; Badre, 2012; O’Reilly & Frank, 2006). This suggests that engaging visual working memory is a distinct process from maintaining additional information in this memory system once it is engaged. One prediction of this account is that when the first item participants see is ‘new’ (not available in visual long-term memory), and thus participants must engage visual working memory to maintain it, they may be more likely to hold the second item in working memory even if it is already available in visual long-term memory. By contrast, if the first item is available in visual long-term memory, working memory may not be engaged at all until the second item is shown.

To address this, we separately analyzed the CDA in the LTM-first and LTM-second conditions and compared them to the LTM-unavailable condition. We entered all three conditions into a repeated-measures ANOVA and observed a significant effect, F(2, 38) = 4.27, p = 0.02, η_p_^2^ = 0.18. Follow up analyses revealed no difference in CDA amplitude between the LTM-unavailable (M = 1.63, SEM = 0.19) and LTM-second (M = 1.58, SEM = 0.30) conditions, t(19) = 0.21, p = 0.84, but a diminished CDA amplitude in the LTM-first condition (M =0.82, SEM = 0.25), which was significantly different compared to the LTM unavailable condition t(19) = 2.71, p = 0.01. Thus, when participants are initially shown an old image during a visual working memory task, followed by a new item, they have the least engagement of visual working memory resources, consistent with the gating hypothesis; whereas when they must engage working memory for the first item if it is new, there is little distinction in working memory activity based on whether the following item is old or new. These effects were not explained by potential differences in attention during the encoding phase, as analyses of P1 and N2PC components (two well-known neural markers of attention; Woodman & Luck, 1999; Moher et al., 2014) found no significant differences across conditions (all p’s > 0.10).

**Figure 3.**
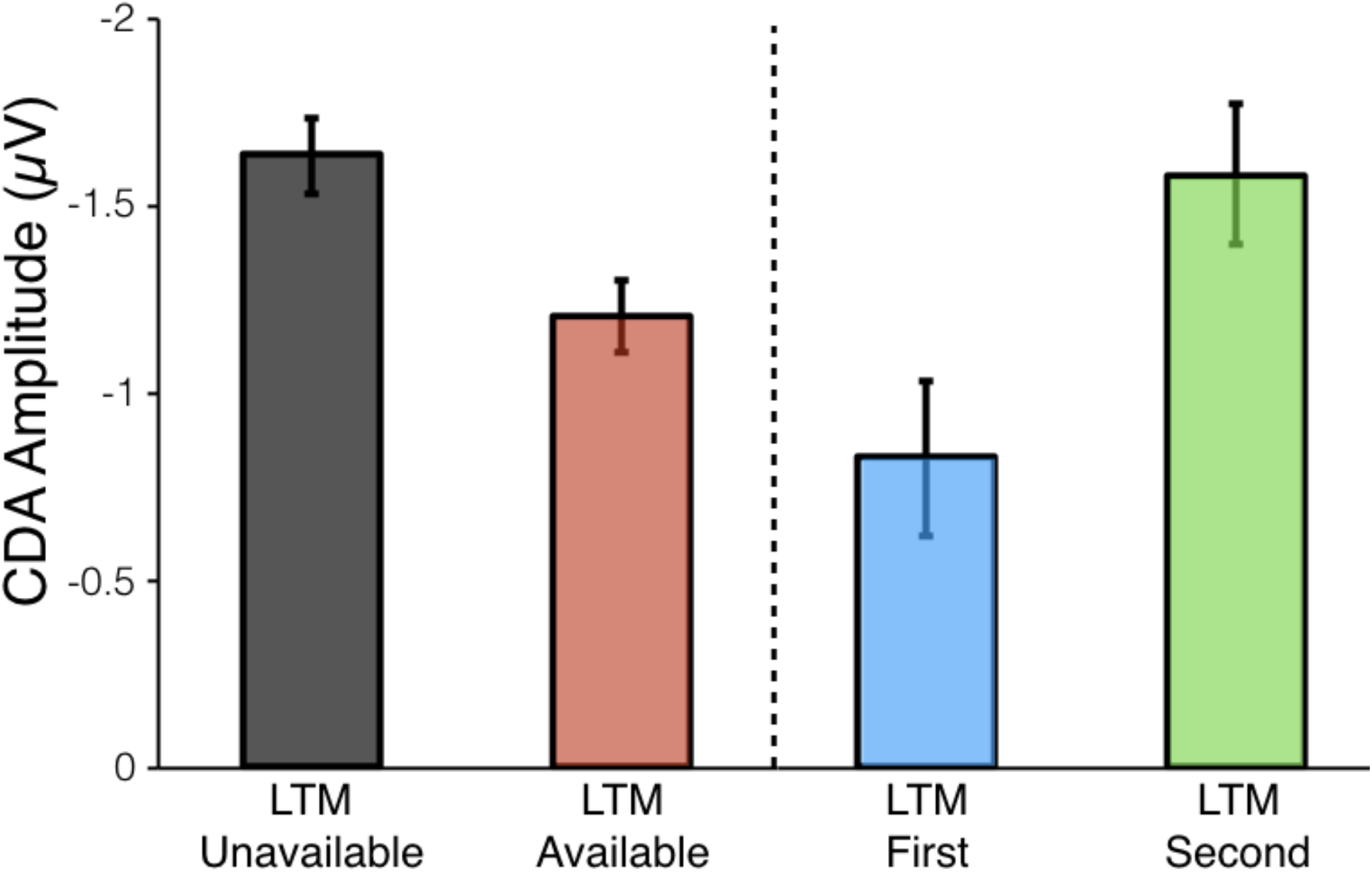
Results of Experiment 1. CDA amplitudes in the LTM-unavailable and -available conditions, as well as the conditions contained within LTM-available (LTM-first, LTM-second). We observed significantly reduced CDA activity in LTM-available compared to LTM-unavailable. Within LTM-available, this reduction was driven by the LTM-first condition: when participants encountered a previously studied item first, the CDA was reduced (as though it was not held in mind actively) but when the previously studied item followed a new item, and so working memory was already engaged, the CDA was not reduced, consistent with the gating hypothesis of working memory. Error bars reflect within-subject standard error.

Could this reduced active maintenance for the LTM-first condition be explained by the idea that working memory resources (reflected by the CDA), regardless of the use of long-term memory, tend to disproportionately allocated to the first item in sequential displays? In theory it is possible that participants could disproportionately allocate their working memory resources to the first item, and then when the first item is new (i.e., LTM-second condition), this results in a larger CDA than when the first item is old (i.e., LTM-first condition). However, existing evidence argues against this idea: in sequential displays, the CDA has been found to more strongly reflect the second set of items rather than the first under some circumstances or approximately equally weigh the two under other circumstances (Berggren & Eimer, 2016; Feldmann-Wüstefeld, Vogel & Awh, 2018). This is the opposite of what we have found in the current study when availability in long-term memory is also manipulated. This suggests that gating provides a better explanation for our data than does a sequential encoding bias.

Broadly, these results suggest that visual working memory, as indexed by CDA activity, is engaged once new information is encountered. However, if a participant first encounters an object previously encoded in visual long-term memory, they maintain significantly less in visual working memory compared to being shown two completely new objects, suggesting they maintain less information about the previously-encoded object in visual working memory and instead rely on their long-term memory representation. Consistent with this, we find that despite maintaining significantly less information in visual working memory as indexed by the CDA, participants do not demonstrate any reduction in behavioral performance in this condition. Overall, the results of Experiment 1 suggest a surprisingly adaptive use of visual working memory only when required, and that participants use visual long-term memory in lieu of visual working memory when it is available. Our results also provide evidence in favor of the ‘gating’ hypothesis – e.g., that engaging working memory may be a distinct process from putting additional information into visual working memory.

### Experiment 2: Working Memory Systems Engage When Perceptual Interference is High

The data from Experiment 1 indicates that participants can dynamically avoid engaging working memory when long-term memory is available, without any detriment to performance. Given these results, what then is the purpose of visual working memory: if long-term memory can be utilized without any apparent costs and with considerably less cognitive demand and capacity limitations, why have a separate, effortful working memory system? Under what circumstances might long-term memory fail to be engaged or utilized over the short-term?

Many researchers have claimed that the main reason we require separate working memory systems is that working memory is less susceptible to inference. For example, consistent with the work of Ericsson and Kintsch (1995), Engle (2002) argued that dealing with interference is the primary function of working memory: "Without the effects of interference, most of the information people know and need to function in the world could be retrieved from long-term memory sufficiently quickly and accurately for them to perform even complex cognitive functions quite well" (Engle, 2002). Thus, the results of our first experiment, where participants quickly and accurately make use of long-term memory instead of engaging visual working memory, may depend crucially on the level of interference participants expect to encounter. To investigate this, in Experiment 2 we introduced substantial perceptual interference into our working memory task by asking participants to remember two objects in memory that were of the same category. The addition of multiple within-category objects to be remembered was designed to decrease the utility of long-term memory; while people have an extremely impressive long-term memory for object details when all objects are unique (e.g., Brady et al. 2008), adding multiple items from the same category can quickly reduce the ability to retrieve perceptual details from long-term memory (e.g., Konkle et al. 2010). For example, Konkle et al. (2010) found that the addition of only 16 items of the same category caused a drop in visual long-term memory performance similar in magnitude to the drop in performance from the addition of 7,500 new categorically distinct items in Standing (1973) and Standing et al. (1970) studies.

After encoding the two objects of the same category, participants then had to make a detailed perceptual comparison involving an object from the previous working memory display and a third similar-looking object. If participants demonstrate no differences in the CDA across conditions, this would suggest that when perceptual interference is high (and thus long-term memory becomes unreliable), participants actively maintain perceptual information in working memory regardless of having a previous representation to rely on (as they do not expect løng-term memory to be sufficient for this task).

## Methods

### Participants

A group of 21 University of California, San Diego undergraduates participated in Experiment 2. The results from one participant were excluded due to technical errors and/or excessive artifacts in the EEG signal, leading to a final sample of 20 participants. All participants reported normal or corrected-to-normal visual acuity. Participation was voluntary, and in exchange for extra credit in related courses. The experimental protocol was approved by the University of California, San Diego IRB.

### Procedure

Experiment 2 was designed to measure whether CDA differences between the LTM-unavailable and LTM-available conditions would persist when substantial perceptual interference was added to the working memory task. Experiment 2 was thus exactly the same as Experiment 1, with the following exceptions:

During each working memory trial, rather than seeing four categorically distinct objects (presented as pairs in each hemifield), objects presented in the same visual hemifield were drawn from the same category (e.g., two teddy bears). This meant that on a given working memory trial, participants were cued to remember two objects from the same category presented sequentially in a particular hemifield. Thus, a separate stimuli database of real-world objects was used which contained three similar items of the same category (Konkle et al., 2010). As in Experiment 1, participants were told to remember the objects on the left or right of fixation and to ignore the other objects. Objects on the to-be-ignored side of the display were randomly drawn from different categories of objects shown on other trials of the tø-be-remembered side. As a result, the visual display on both hemifields were equated for perceptual information so that any brain differences between the cued and uncued side during the delay were due to differences in working memory, and not perceptual processing. Consistent with Experiment 1, on half of the trials both of the to-be-remembered objects presented were new. For the other half of the trials, one of the objects encountered had been previously encoded during the long-term memory encoding session. Of these trials, half of the time the long-term memory item was encountered as the first item in the trial sequence (“LTM-first”), whereas for the other half the long-term memory item was encountered as the second item in the sequence (“LTM-second”).

At subsequent test, participants were presented a perceptual discrimination forced-choice test and remained visible until participants made a response. This test consisted of one of the two previously studied items from the working memory display (counterbalanced equally across all conditions) and a third, similar lure from the same category, with both items appearing above or below fixation on the to-be-remembered side.

**Figure 4.**
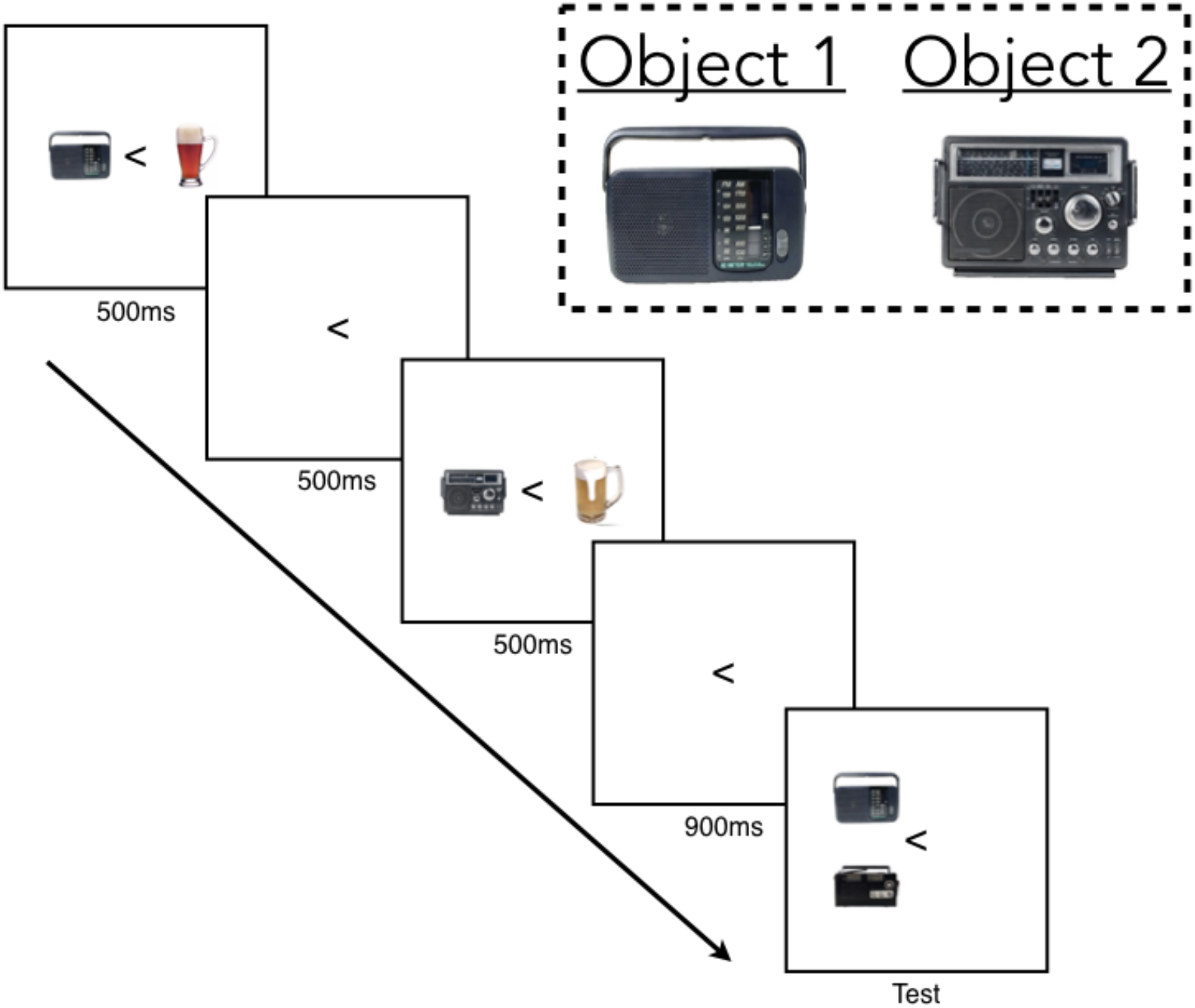
Design of the sequential working memory task used in Experiment 2. The task was the same as Experiment 1, except for the addition of substantial perceptual interference through having participants remember two objects from the same category during encoding. At test, observers saw one of the two previous objects above or below fixation, paired with a third new object (lure) from the same object category. The addition of multiple within-category objects to be remembered was designed to decrease the utility of long-term memory; while people have an extremely impressive long-term memory for object details when all objects are unique (e.g., Brady et al. 2008), adding multiple items from the same category can quickly reduce the ability to retrieve perceptual details from long-term memory (e.g., Konkle et al. 2010).

### Electrophysiological Recordings and Analysis

EEG recordings and analysis were the same as in Experiment 1, with the exception that in Experiment 2 (collected at University of California, San Diego), data were online referenced to the right mastoid and later were offline referenced to the average of the right and left mastoid.

## Results

Consistent with Experiment 1, we again found that across both main conditions accuracy was similar. In particular, participants averaged 88.2% correct (SEM = 0.8%) in the LTM-unavailable condition and 88.6% correct (SEM = 1.0%) in the LTM-available condition, which were not significantly different from one another, t(19) = 0.92, p = 0.37, Cohen’s dz = 0.12. However, we did observe an effect of order in the LTM-available condition. Participants averaged 90.9% correct (SEM = 1.0%) in the LTM-first condition and 86.3% correct (SEM = 1.4%) in the LTM-second condition, t(19) = 3.84, p = 0.001, Cohen’s dz = 0.85. The significant effect was primarily driven by better performance for when the first item was tested in the LTM-first compared to LTM-unavailable condition (M = 90.7% vs 85.8%, respectively), t(19) = 2.76, p = 0.01, Cohen’s dz = 0.62. We failed to find any other significant differences for all remaining contrasts (all p’s>0.20). Thus, it appears that in the LTM-first condition, observers were able to use previously encountered memories for objects to gain a slight performance advantage under very specific conditions (i.e. only for the first item). Note that because we needed to use a different stimulus set in Experiment 2, performance here is not directly comparable to performance in Experiments 1.

**Figure 5.**
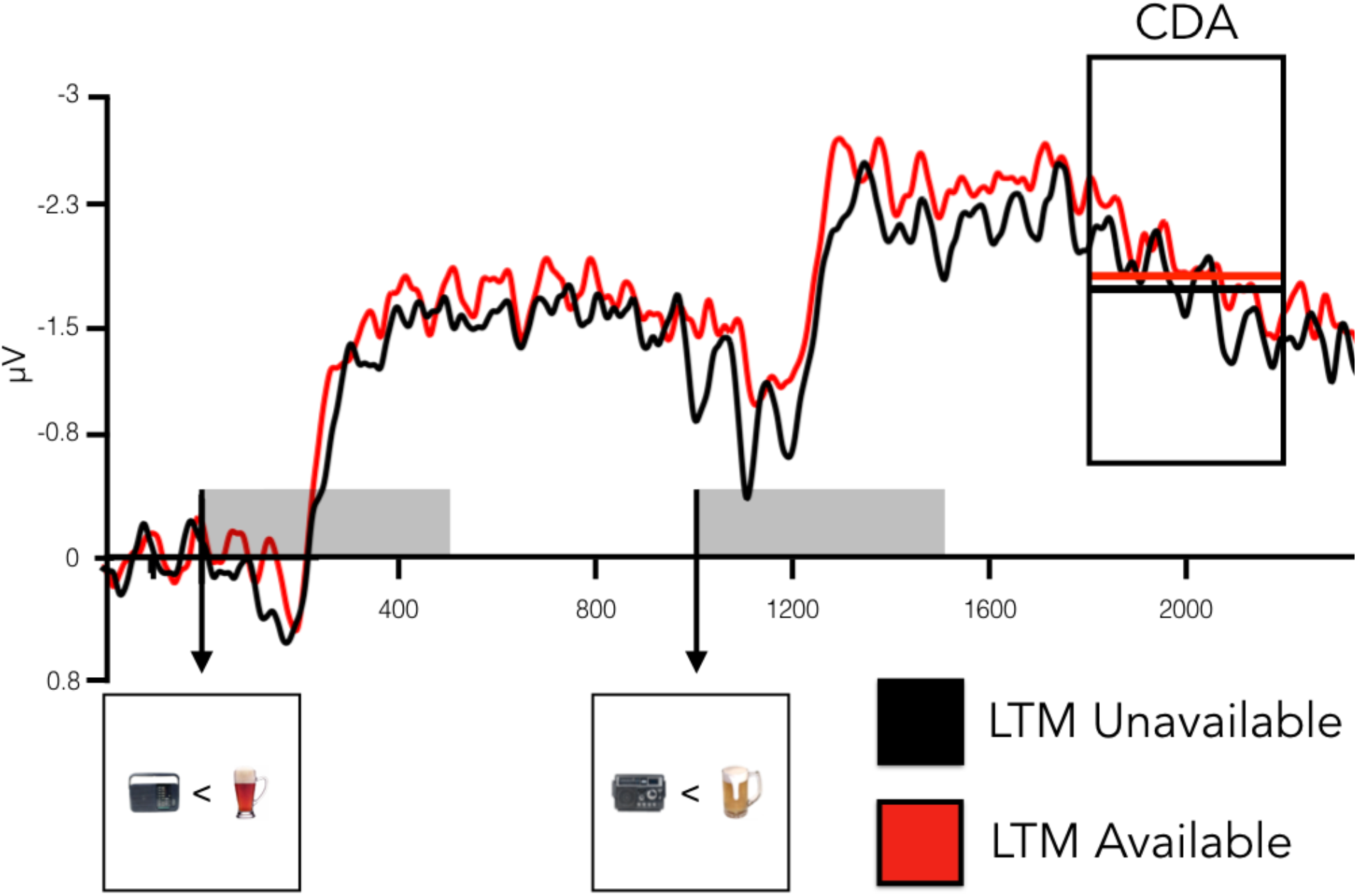
Results of Experiment 2. Contralateral-minus-ipsilateral waveforms for the LTM-unavailable (black) and LTM-available (red) conditions. The CDA is measured from 300 ms after offset during the delay period (black rectangle, labeled CDA). When substantial perceptual interference was introduced into the working memory task, we failed to find any differences between the CDA amplitudes for when participants had a LTM representation available, compared to when they did not.

We were primarily interested in whether we would observe any differences in CDA activity across conditions when perceptual interference was high. First, we compared trials containing only new images (LTM-unavailable) to trials containing a previously seen long-term memory image (LTM-available). CDA amplitude was not significantly different according to whether one object had been previously encountered in visual long-term memory (M = 1.91, SEM = 0.30) compared to when both images were new (M = 1.72, SEM = 0.35), t(19) = 0.95, p = 0.35. Next, all three conditions were entered into a repeated-measures ANOVA. We failed to observe any significant differences, F(2, 38) = 0.54, p = 0.59, η_p_^2^ = 0.03. Therefore, CDA amplitude was not statistically distinguishable between the LTM-unavailable, LTM-second (M = 1.86, SEM = 0.32) and LTM-first (M = 1.97, SEM = 0.33) conditions (p>0.20). This suggests that under conditions where participants expected to encounter sufficient interference to decrease the utility of their long-term memories (Konkle et al. 2010), participants actively maintained the same amount of information in all conditions, regardless of whether or not they had a previous long-term memory representation available. Thus, during conditions of low expected interference (Exp. 1) participants quickly and easily replaced maintenance in visual working memory with access to long-term memory, whereas under conditions of high interference (Exp. 2) they engaged working memory regardless of whether they had long-term memories available. This provides evidence consistent with the idea that one of the main purposes of working memory broadly, and visual working memory specifically, is to provide a memory system that is robust to interference.

**Figure 6.**
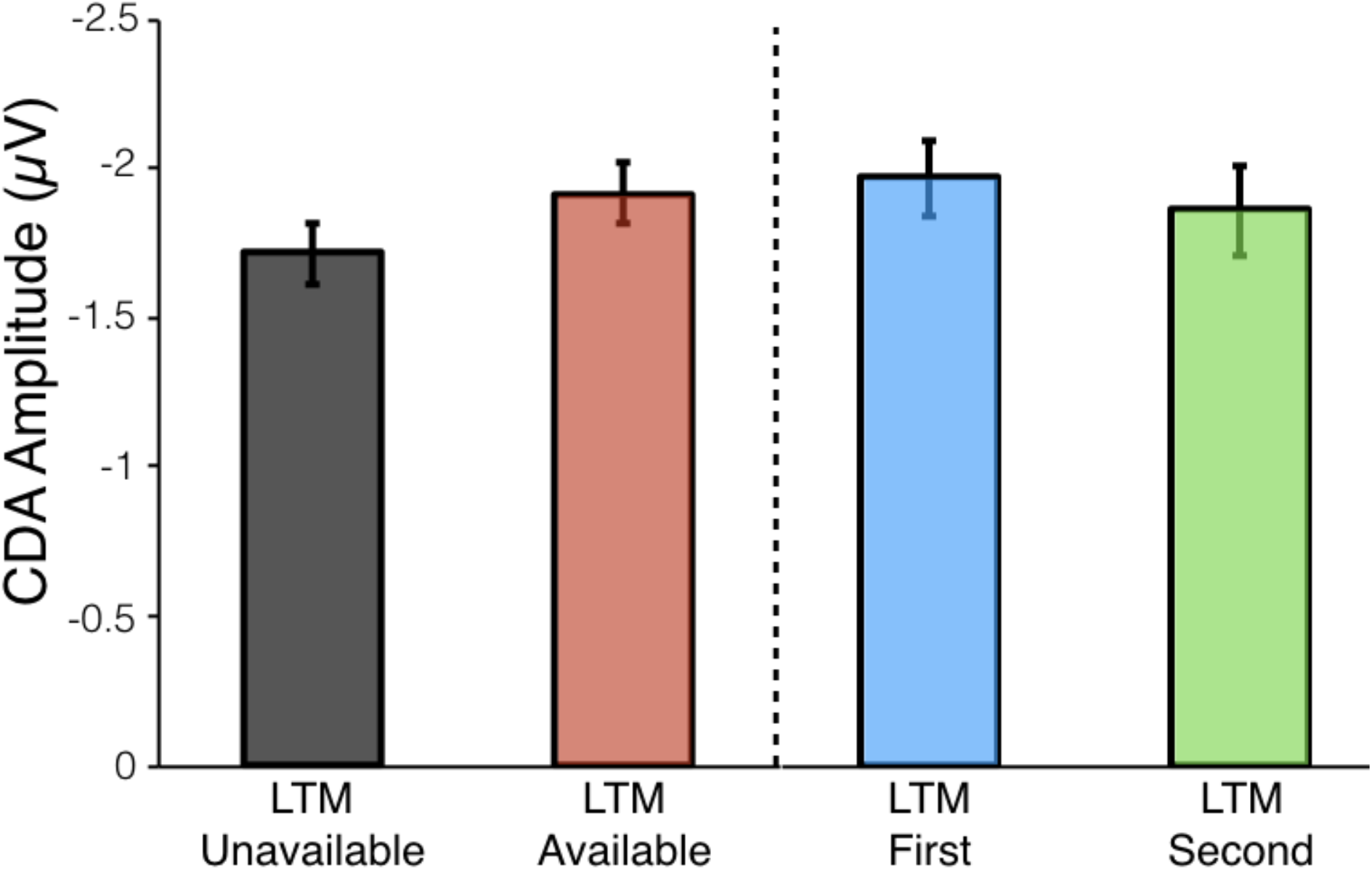
Results of Experiment 2. CDA amplitudes in the LTM-unavailable and -available conditions, as well as the conditions contained within LTM-available (LTM-first, LTM-second). Under conditions of increased perceptual interference, we failed to find any difference across conditions in active maintenance in working memory (as indexed by the CDA). Error bars represent within-subject standard error.

## Discussion

The present results show that under conditions of low interference, having long-term memory representations available significantly reduces participants’ use of active maintenance in visual working memory. Specifically, we find that working memory is engaged once new information is encountered or when perceptual interference is high. However, when interference is low and participants have a previous visual long-term memory to rely on, this existing visual long-term memory can be utilized in a short-term memory test instead of engaging working memory systems. Critically, these long-term memory representations contain enough visual details (see Brady et al., 2008) to be used without impacting performance, even when making fine perceptual discriminations at test, as long as interference is low. Thus, the present data reveals how visual long-term memory representations may be used to support memory over brief durations. Overall, this suggests a more intimate and dynamic interaction between working memory and løng-term memory than is typically considered, with visual long-term memories of previous encounters with objects substituting for working memory in a setting that is typical of a real-world task (e.g., where items are not novel and meaningless). The limited capacity of visual working memory may therefore be less of a constraint on performance than is typically assumed, if when performing real-world tasks we are often able to make use of our impressive visual long-term memory capacities rather than holding items actively in visual working memory.

### Potential dynamics between long-term and working memory

While we show that participants can use long-term memory in lieu of active maintenance in working memory, it is also likely that under some circumstances participants could be incentivized to use both memory systems, resulting in a performance advantage when LTM is available but no change in how much is stored in working memory (e.g., no change in CDA). That is, working memory engagement is generally under active conscious control (e.g., Baddeley, 2003). We are not claiming that long-term memory must always replace active maintenance in working memory, only providing a demonstration that such replacement is possible under realistic conditions, and thus that in real-world scenarios participants may often be able to make use of visual long-term memory rather than holding items actively in visual working memory.

An important aspect of our finding of reduced engagement of working memory is that we are using the CDA as our measure of working memory, which indexes only perceptual working memory for visual information, not all forms of what might be considered ‘working memories’. Thus, our finding of reduced CDA could arise from a long-term memory representation becoming active again during the working memory task, but with the format of that representation no longer as perceptual as an object that enters working memory directly from perceptual experience. For example, the format of the working memory trace might be some code, label, or placeholder which is an efficient representation that allows participants to access the long-term memory (e.g., Ericsson & Kintsch, 1995). From this perspective, the long-term memory objects might still be maintained in working memory but in a format that isn’t reflected by the CDA. In the present task, observers could not have relied on a totally abstract working memory trace, such as a label or placeholder, given they needed to complete a perceptual discrimination at test (which required visual details). However, it still remains possible that when recognizing a long-term memory item, either a pointer to the long-term memory trace is maintained in working memory or the details of that item are maintained, but in a different format not reflected by activity in the CDA. This would still suggest that participants can recognize the existence of a previous memory trace within a few hundred milliseconds, and adapt their encoding strategy to depend on this memory trace, all for an object they have only encountered briefly one time. However, this account would suggest that the existence of a løng-term memory affects working memory not by reducing perceptual maintenance per se but rather through altering the format of the working memory representation itself.

In addition, it is important to note that we are referring to working memory usage during the delay period, and that at subsequent test it remains unclear how long-term memory representations may be utilized in conjunction with working memory resources. For example, it could be that when a long-term memory item is tested, perceptual information is pulled from long-term memory and held in working memory in order to make a response (Fukuda & Vogel, 2017). In addition, it is important to emphasize that the ability of working and long-term memory to trade-off and interact with one another is likely constrained by specific circumstances. While in Experiment 1 we demonstrated reduced perceptual maintenance for items with previous long-term memory representations, in Experiment 2 we were able to eliminate this effect by substantially increasing the amount of perceptual interference in the working memory task. Indeed, perceptual interference is likely only one of what are several factors that may reduce or eliminate the ability of long-term memory to interact with and potentially support working memory. It remains an interesting venue for future research to further clarify the conditions under which working and long-term memory interact (or fail to interact) with one another.

### Possible mechanisms supporting dynamic utilization of long-term memory representations

Initially, the flexibility and speed with which long-term memory representations can be utilized in a short-term memory task appears quite surprising. For example, in the LTM-first condition, there must be a process that allows the image to be recognized quickly enough that working memory for that item is not engaged. What are the potential mechanisms that may facilitate this process?

There is significant evidence, for example from Ericsson and Kintsch (1995), that participants are able to quickly and effectively access long-term memory systems to perform tasks. This may be especially true when all that is required is familiarity (that is, no source memory is required). Consistent with this idea that after encoding dozens of images into visual long-term memory participants may have fast and efficient access to whether a given image is familiar or not, research into hybrid visual search has shown participants can hold hundreds of potential targets in long-term memory and efficiently search for any of these items (Wolfe, 2012; Cunningham & Wolfe, 2014).

Could it also be possible that participants are using long-term memory, but relying on a recollection signal as opposed to a familiarity-based signal? Research suggests that participants likely could not rely on recollection. In work conducted by Carlisle et al. (2011), observers were given an attentional task where only a few very simple stimuli needed to be held in mind (as search targets), and thus people had long-term memories for each item. Despite substantial experience with a small number of items, they observed participants still demonstrated a large CDA the first time they had to use one of those items as a target, even when it had been a target before. However, the CDA did decrease over repetitions of trials in a row that had the same item as a target (Carlisle et al., 2011). This suggests that for simple stimuli people cannot instantly access whether it is an item they have in long-term memory, and must therefore rely on working memory. Only after several repetitions of the same target – by which time it is by far the most familiar of the stimuli again – can participants easily recognize the item without working memory.

Previous research has also demonstrated familiarity signals for items occur quite rapidly (within 100 ms), much faster than recollection signals required for source memory (at least 500 ms) (Hintzman, Caulton & Levitin, 1998). Similarly, novelty signals – which may be used in order to assess whether an item may require the engagement of working memory – are also extremely fast. For example, previous research in monkeys has shown visual novelty can trigger neuronal population firing 70-80 ms after stimulus onset (Li, Miller & Desimone, 1993; Xiang & Brown, 1998; Brown, 2009). In the context of the present task, during a trial of the working memory task, familiarity or novelty is all that is required for an observer to know they can rely on a løng-term memory representation during the subsequent test. Thus, even our brief encoding times for individual images (500 ms) are more than enough time for a familiarity or novelty signal to prevent working memory from engaging. It remains an open question whether the same effects would be observed if recollection was required to recognize objects in working memory, and not just familiarity. Future research could disentangle the dynamics and systems supporting the trade-off between working and long-term memory, with conditions that only require familiarity (as in the present series of experiments) compared to others that require recollection to recognize objects when presented in working memory.

Importantly, this fast access to items in long-term memory may not occur for all previously encoded items, but may require those memories to be held in a special state that allows for fast access, sometimes referred to as activated løng-term memory (e.g., Cowan, 2008). In the hybrid search literature, there is some evidence that items that are expected to be used in the task are more accessible or active than other items (e.g., Boettcher, Drew, & Wolfe, 2013). The extent to which items need to be held in an activated long-term memory state to replace working memories remains an open question.

### Support for gating hypothesis of working memory

In Experiment 1, we observed that CDA amplitude in the LTM-second condition was analogous to activity observed during LTM-unavailable trials. This is consistent with the “gating” hypothesis of working memory. This theory suggests that in order to balance flexibility and stability, working memory operates via separate gate and maintenance mechanisms: when the gate is open available information can enter working memory, and when the gate is closed the contents of working memory are sustained while keeping irrelevant information out (Miller, Cohen, 2001; Raghavachari et al., 2001; Badre, 2012; O’Reilly & Frank, 2006). Separating maintenance and gating into distinct mechanisms has been shown to be computationally efficient (Hochreiter & Schmidhuber, 1997) and remains an assumption in several prominent and influential working memory models (Braver & Cohen, 2000; O’Reilly & Frank, 2006).

Thus, in the context of the present studies, when the first item presented in a working memory display was one encoded previously in visual long-term memory, observers were able to utilize long-term memory representations as the working memory gate had not been opened. But when the first item encountered was new and had not been previously encoded, the working memory gate was opened to maintain subsequently encountered information. As a result, in the LTM-second condition, even though observers had a long-term memory available for the second item encountered, they encoded this information into working memory.

## Conclusions

Our data illuminate the conditions under which long-term memory and working memory operate. Working memory is engaged when new, previously unstudied information is encountered or when perceptual interference is high. In contrast, when old information is present or perceptual interference is low, participants may use long-term memory in lieu of working memory. These results further our understanding of how and when hand-offs occur between long-term memory and working memory systems. Moreover, our findings demonstrate not only how these systems interact with one another, but also suggest the relationship between working memory and løng-term memory is more dynamic and fluid than previously thought.

## Acknowledgements

Some of the results were presented at the annual meeting on Object Perception, Attention and Memory (OPAM) in 2017 and the annual meeting of the Vision Sciences Society (VSS) in 2018. For helpful feedback on this work, we would like to thank Viola Störmer and John Wixted.

